# MoMo - Combining Neuron Morphology and Connectivity for Interactive Motif Analysis in Connectomes

**DOI:** 10.1101/2025.07.02.662847

**Authors:** M. Shewarega, J. Troidl, O. Alvarado Rodriguez, M. Dindoost, P. Harth, H. Haberkern, J. Stegmaier, D. Bader, H. Pfister

## Abstract

Connectomics, a subfield of neuroscience, reconstructs structural and functional brain maps at synapse-level resolution. These complex spatial maps consist of tree-like neurons interconnected by synapses. Motif analysis is a widely used method for identifying recurring subgraph patterns in connectomes. These motifs, thus, potentially represent fundamental units of information processing. However, existing computational tools often oversimplify neurons as mere nodes in a graph, disregarding their intricate morphologies. In this paper, we introduce *MoMo*, a novel interactive visualization framework for analyzing *neuron morphology-aware motifs* in large connectome graphs. First, we propose an advanced graph data structure that integrates both neuronal morphology and synaptic connectivity. This enables highly efficient, parallel subgraph isomorphism searches, allowing for interactive morphological motif queries. Second, we develop a sketch-based interface that facilitates the intuitive exploration of morphology-based motifs within our new data structure. Users can conduct interactive motif searches on state-of-the-art connectomes and visualize results as interactive 3D renderings. We present a detailed goal and task analysis for motif exploration in connectomes, incorporating neuron morphology. Finally, we evaluate *MoMo* through case studies with four domain experts, who asses the tool’s usefulness and effectiveness in motif exploration, and relevance to real-world neuroscience research. The source code for *MoMo* is available [here](https://github.com/VCG/momo).

## 1 Introduction

Connectomics is a rapidly advancing subfield of neuroscience focused on mapping the intricate network of connections between neurons down to the level of individual synapses. The overarching goal is constructing a comprehensive wiring diagram of an organism’s nervous system. Recent breakthroughs [17, 30, 56, 64] in imaging and automated neuron reconstruction have made this vision increasingly tangible. This resulted in the publication of a complete connectome of the fruit fly brain [16] as well as cubic-millimeter-scale connectomes of both mouse [65] and human [60] brain tissue. These datasets now enable scientists to trace complete neural circuits that underlie all aspects of information processing in the brain. Analyzing such circuits has already yielded transformative insights into the neural basis of behavior. For instance, studies of the fly connectome have advanced our understanding of navigation [29], vision [47], and motor control [36], among many others. However, extracting meaningful information from large-scale connectomes remains an enormous challenge due to several key factors. First, modern connectomes are extraordinarily large, exceeding a petabyte size, making them computationally intensive to process. Second, neurons are complex, tree-like structures that extend across significant spatial distances. This morphology complicates the interpretation and querying of connectivity. Third, the connectivity graphs are extremely dense: a single neuron can form synapses with over 1,000 partners, rendering traditional node-link diagrams ineffective for visual analysis. To address this complexity, neuroscientists often employ motif analysis [62] as a divide-and-conquer strategy, breaking down large connectomes into smaller, more interpretable subgraphs. For example, motifs in the H01 dataset [60] have been identified based on strong pairwise connections via multiple nearby synapses. Similarly, motifs involving compass neurons in the fruit fly’s central brain have elucidated mechanisms for encoding directional information [29]. Crucially, these analyses consider both synaptic connectivity and neuron morphology. Nonetheless, current interactive tools for connectomic analysis [39, 40, 52, 68] lack support for querying networks based on both synaptic connectivity and morphological structure.

In this paper, we present *MoMo*, a novel interactive visualization and analysis tool designed to enable neuron morphology-aware motif analysis in large-scale connectome graphs. Our contributions are four-fold: **(1)** First, we conduct a detailed study of domain-specific goals for interactive motif analysis incorporating neuron morphology. Based on these insights, we derive a set of analytical tasks that *MoMo* must support to meet neuroscientists’ needs. **(2)** Second, we introduce a novel two-layer graph-based representation for connectomes that jointly encodes neuronal morphology and synapse-level connectivity (Sec. 6). At the morphology layer, each neuron is modeled as a set of interconnected segments capturing its branching structure. At the connectivity layer, synaptic edges link segments across neurons. This separation of morphology and connectivity reduces data size compared to voxel-based representations and enables the application of established graph analysis techniques. **(3)** Third, we implement a Jupyter-based prototype of *MoMo* that supports the full set of analysis tasks. The tool features an intuitive sketching interface that allows users to draw motifs—including both morphological components and synaptic connections—as query patterns. These sketches are used to search the underlying graph representation, and matching instances are visualized in interactive 3D renderings, allowing detailed inspection of how motifs are embedded within neural tissue. **(4)** Fourth, we evaluate *MoMo* through pilot and case studies on two state-of-the-art connectome datasets, conducted in collaboration with expert neuroscientists. Each neuroscientist selected a phenomenon of interest, such as *shunting*/*lateral inhibition* or *center-surround receptive fields*, and successfully sketched and queried for corresponding motifs. *MoMo* enabled experts to analyze potential instances of these phenomena by sketching individual neuron segments and synapses—granularity not supported by existing cell-level abstractions (see Section 10 for examples).

## 2 Related Work

### Visualizing Connectomes

Beyer et al. [8] provide a comprehensive survey of interactive visualization techniques for connectome analysis. Prior work can be broadly categorized into: (a) data structures and algorithms for interactive analysis [6, 7, 23, 25, 61], (b) interactive spatial exploration [5,26,45,55,66], (c) connectivity analysis [1,21,22,55,68–70], and (d) visualization for scientific communication [11,12]. Ganglberger et al. [23] introduce a spatially-driven visual analytics framework for iterative exploration of large, heterogeneous brain datasets; however, it lacks fine-grained motif search at the level of individual neuron segments and synapses. *MoMo* advances the state of the art by integrating interactive spatial analysis, connectivity analysis, and connectomic data representation. Unlike previous systems that typically explore morphology and connectivity separately [68], *MoMo* supports integrated, interactive queries combining both via a novel graph-based representation derived directly from neuronal skeletons, enabling morphology-aware motif searches. Existing methods often simplify the connectome into node-link diagrams, with nodes as entire neurons and edges as synapses. While dendrogram-based approaches [63] capture some neuronal structure, they do not support flexible, targeted motif queries. By bridging structural morphology and connectivity, *MoMo* enables new interactive analysis modes for large-scale connectomic datasets.

### Visual Motif Analysis

Visualizing network motifs is a well-established task in network visualization [10, 19, 34], with conventional techniques primarily applied to domains such as social networks, citation graphs, and web connectivity. However, these approaches do not directly extend to connectomic data due to the inherently three-dimensional spatial structure of neurons—the nodes of connectome graphs—which introduces domain-specific challenges for motif visualization. To address this, Vimo [68] introduced an interactive system for constructing and querying motifs within connectomes. *MoMo* builds on this foundation by enabling the specification and analysis of morphological motifs—substructures that incorporate both neuronal branching geometry and precise synaptic connectivity. This extension allows users to define queries that account for spatially and structurally localized connectivity patterns, a critical feature for understanding neural circuits. In parallel, computational methods have been proposed for motif detection in connectomics. Matejek et al. [38] presented a parallelized algorithm for efficient motif enumeration in large-scale brain datasets. At the same time, DotMotif [40] introduced a domain-specific language designed to express motif queries in a concise, readable form, with backend integration into Python and Cypher [20]. While these systems focus on computational efficiency and query abstraction, *MoMo* complements them by providing an interactive, visual environment tailored to spatially embedded morphological motifs in connectome data.

### Visual Graph Query Interfaces

Interactive visual graph query interfaces have been explored in various domains, such as bibliographic data [74], and genomics data [43], among many others. Visage [50, 51] shows how visual graph query interfaces can simplify pattern analysis in simple conventional graphs, such as movie and actor networks derived from Rotten Tomatoes. Follow-up work [49] has investigated new visualization approaches to inspect and analyze motif query results in conventional graphs. However, those previous approaches do not translate directly to morphological brain motif analysis since these approaches are designed for data without three-dimensional spatial context. VisualNeo [28] aims to bridge the design for graph query engines and visual graph query interfaces through a custom-designed software system. VIIQ [31] simplifies interactive query construction using an edge suggestion algorithm. Existing visual query systems for spatio-temporal data, such as Diehl et al. [14], focus on interactive filtering and exploration of weather forecast data and include a curve-pattern selector for specifying and searching temporal motifs. However, their system does not support the definition or querying of branching or connectivity patterns, nor does it provide general sketch-based motif specification for spatially embedded structures.

### Multi-layer & Micro-vascular networks

McGee et al. [42] survey the state of the art in multi-layer network visualization. Most relevant to our work are multiplex network visualizations [9, 46], which are defined by the presence of various edge types. However, most use cases center around applications in sociology. Most similar to our work is the visualization of vascular networks [24]. For example, Mayerich et al. [41] visualize volumetric vascular data using techniques from volume rendering and do not explicitly extract the graph information.

### Previous Connectivity Motif Workflow

Vimo [68] offers a user-friendly interface that allows users to draw motifs using a data abstraction where each node represents an entire neuron and each edge represents a synaptic connection between neurons. While this abstraction simplifies the visualization and allows for an overview of which neurons are connected, it has significant limitations. One major limitation of Vimo’s approach is its lack of detailed morphology information. In Vimo, each neuron is represented as a single node, which means the internal structure—such as the exact locations of synapses, branching points, and neuron morphology—is completely lost. This makes it impossible to perform fine-grained analyses that require insight into the internal pathways and synaptic organization within a neuron.

## 3 Biological Background

### Neuroscience Fundamentals

Brain tissue consists of neurons, among other cells, that are connected through synapses [27] (see Fig. 2). Neurons are complex, tree-like spatial structures that connect in various spatial configurations. The spatial configuration of neuron connectivity can heavily influence the underlying function of the respective neuronal circuit. The resulting network diagrams are exceptionally dense, with current connectome datasets exceeding tens of thousands of neurons interconnected through millions of synapses.

**Fig. 1:**
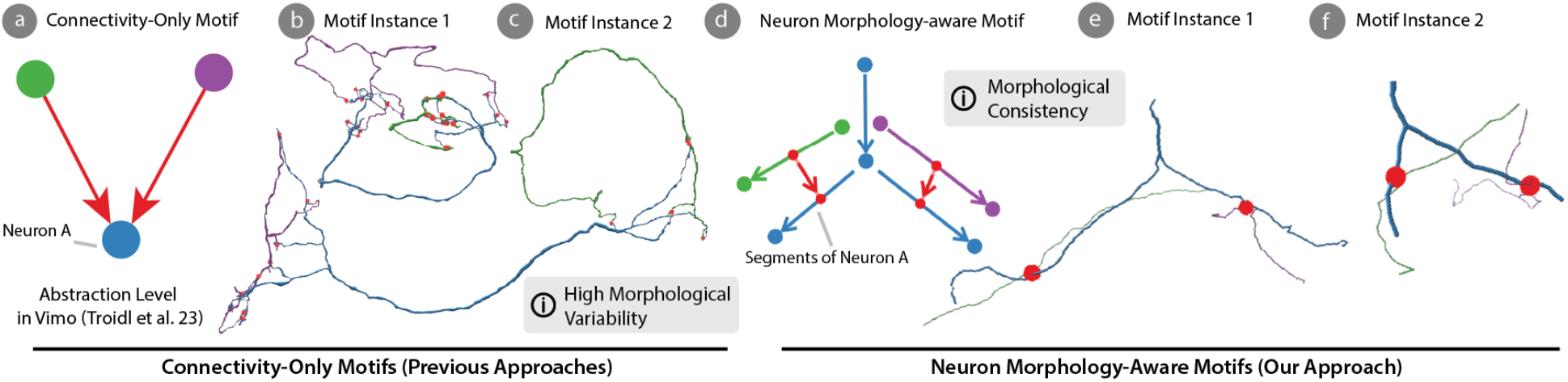
Connectivity-only vs. Neuron Morphology-Aware Motif Analysis. (a) Previous work [40, 68] has focused on interactive motif analysis, where neurons are represented as simple nodes connected through synapses. (b, c) This representation lacks scientifically relevant neuron morphology, leading to ambiguity in motif queries. (d) Our approach integrates neuron morphology into motif queries and enables searching motifs interactively by explicitly sketching neuron segments and their respective synapses. (e, f) Hence, users can perform targeted queries for both morphological and connectivity patterns. Data: FlyEM Hemibrain (a-c) [57], MICrONS (d-f) [65].

**Fig. 2:**
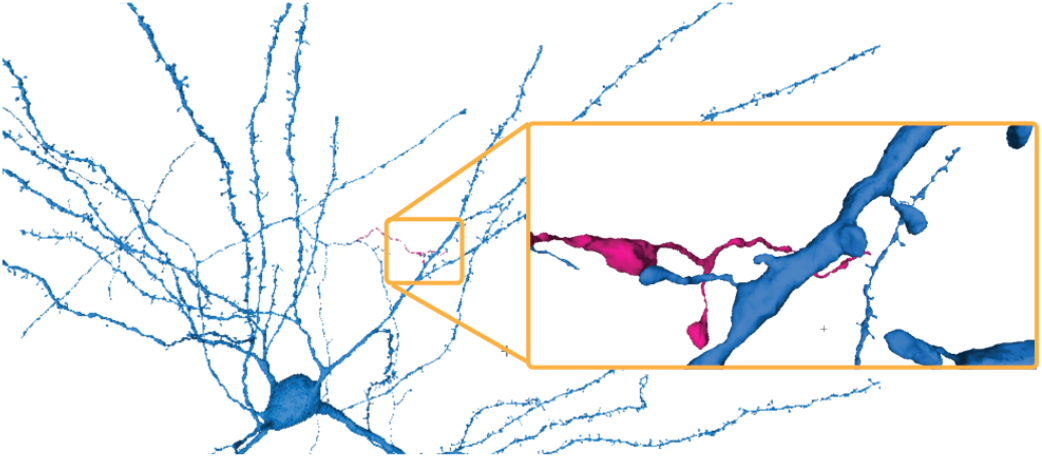
Data Overview. Modern connectomics reconstructs tens of thousands of neurons connected through millions of synapses. Here, we show a single neuron (blue) and a single synaptic connection to a pink neuron (orange box). Data: Human cerebral cortex / H01 [60].

### Connectomics Data

Creating connectomes is an involved process that starts with acquiring 3D image volumes, typically using electron [56] or optical [64] microscopes. The resulting volumes are then segmented using 3D convolutional neural networks [30]. Next, the automatically reconstructed neurons and synapses undergo human or automated proofreading [17, 33, 67, 72], eliminating errors from the previous steps. Here, we demonstrate our approach on two exemplary state-of-the-art connectomes. We use the *FlyWire* connectome [16] for insect neurons and the *MICrONS* dataset [65], which reconstructs approximately one cubic millimeter of mouse visual cortex. For both datasets, *MoMo* uses neuron skeletons and synaptic point annotations.

### Motif Analysis in Connectomes

Neuronal connectivity motifs are fundamental building blocks of computation in the brain. Many neuroscientific studies [16, 29, 60] are searching and analyzing structural and connectivity motifs in the brain. For example, Hulse et al. [29] report a group of neurons in the fruit fly’s central brain that form motifs, which allow flexible spatial navigation and action selection. Another exemplary motif that combines specific connectivity and neuron morphology patterns is the concept of *shunting inhibition* [32,44]. Shunting inhibition is a fundamental mechanism in neural computation, where inhibitory synapses modulate the excitability of a neuron by altering its input resistance. This mechanism effectively “shunts” incoming excitatory signals, thereby regulating synaptic integration and influencing the neuron’s ability to fire an action potential. Shunting inhibition is crucial in various neural processes such as sensory processing or network synchronization, among others. We report an example of *shunting inhibition* in the case study (see Sec. 10).

## 4 Goals and Task Analysis

The idea of integrating neuron morphology into tools for motif analysis originated while struggling to represent biologically relevant motifs [60] during the development of Vimo [68]. As a result, we discussed extending Vimo to morphological motifs in informal interviews with four domain experts at an international connectomics conference and research visits at both the Harvard Center for Brain Science and Howard Hughes Medical Institute (HHMI) Janelia. All scientists are leading experts in analyzing neuronal circuits reconstructed from high-resolution electron microscopy data. Two experts focus on analyzing mammalian brain tissue, while the others specialize in the fruit fly (*Drosophila*) brain. Following the design study methodology from Sedlmair et al. [59], we distilled the following goals and tasks.

### 4.1 Domain Goals

#### G1 - Quick Identification of Motifs

Hypothesis generation and exploratory analysis are critical while analyzing connectome circuits. Additionally, data exploration may generate new hypotheses, which in turn requires rapidly iterating specific motifs of interests. Thus, domain experts need tools to quickly identify various motifs of interest.

#### G2 - Precise Connectivity and Morphology Searches

The function of neuronal circuits is governed by both neuron morphology and synaptic connectivity [29, 60]. Queries such as *“Which neurons have multisynaptic connections on the same neuronal branch?”* or *“Which neuron pairs have connectivity clusters on separate branches?”* are common questions in connectome analysis. Thus, neuroscientists need tools to find morphological motifs in conjunction with network motifs. **G3 - Accurate Spatial Analysis of Motif Instances**. After identifying motifs involving neuron morphology and synaptic connectivity, domain experts want to inspect spatial models of the respective neurons and their connectivity. When inspecting neurons and their connectivity, neuroscientists need to (a) view accurate 3D models while (b) correlating the motif query to the original neuronal data.

#### G4 - Flexibility & Data Adaptability

Connectome analysis is a fast-paced scientific discipline with dozens of datasets published annually. Datasets are typically made accessible to the neuroscience community through various platforms [17, 35, 52], which offer simple pythonic interfaces for targeted data retrieval. Thus, scientists need flexible tools that integrate into established, notebook-based analysis workflows without building custom data interfaces.

### 4.2 Tasks

#### T1 - Fast Morphological Motif Queries

Scientists need to define motif queries rapidly but also require computationally efficient subgraph isomorphism searches to identify motifs of interest quickly. (**G1**)

#### T2 - Interactive Sketching-based Queries

Scientists need to precisely define neuron morphology and synaptic connectivity patterns without learning complex graph query languages like Cypher [20] or Gremlin [53]. Interactive sketching-based drawing interfaces are expressive but do not require extensive graph query coding skills to define morphology-aware motifs. (**G1, G2**)

#### T3 - Correlating Motif Structure with 3D Renderings

After a successful query, users must identify the motif structure in a detailed 3D rendering of the respective neurons. (**G3**)

#### T4 - Analyze Motifs in Hyrid Code-UI Workflow

The computational Jupyter Notebook environment brings ease of Python-based data handling and interactive visualization components closer together. Here, data can be loaded directly from various hosting platforms into respective interactive visualization widgets. (**G4**)

## 5 *MoMo* Design and Workflow

*MoMo* provides an intuitive platform for visualizing and analyzing motifs in connectome data that integrates into the computational analysis workflow of connectomics scientists, which often involves data analysis scripts in Python-based Jupyter Notebooks. Jupyter Notebooks provide a robust and interactive environment for quick experimentation and effective visualization. Hence, we designed *MoMo* as a Python package to launch interactive Jupyter widgets for visualization. Users are empowered to programmatically manipulate data before feeding it into *MoMo* but can still use interactive visualization through custom widgets **(G4)**. We designed a simple workflow for *MoMo* (see Fig. 3): **Data Transformation (Sec. 6.2)**. The first step involves transforming the desired connectome dataset into the *neuron morphology-aware data representation*, which enables efficient motif queries (**G2**).

**Fig. 3:**
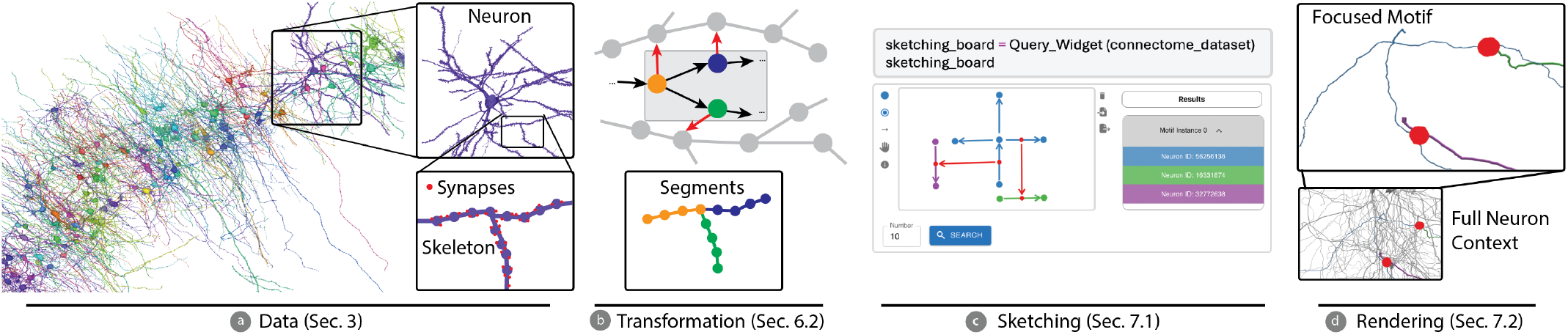
*MoMo* Workflow. (a) Given a connectome dataset, we (b) transform the respective neurons and synapses into our neuron morphology-aware graph representation. Each node in the graph depicts a neuron segment that is connected to other segments through synaptic connections (red arrows) or to neighboring segments (black arrows). (c) The transformed dataset is interactivley queried by sketching motifs. (d) Finally, users can explore identified motif instances in interactive 3D renderings and highlight the motif’s morphology through a focus & context approach. Data: Human temporal cortex / H01 [60] (a), MICrONS [4] (d).

**Sketching (Sec. 7.1)**. Users interactively draw a motif at the neuron segment level using the *sketching interface* (see Fig. 6) **(G1, G2)**. After the user defines a motif, *MoMo* enables highly performant isomorphic subgraph queries (Sec. 8) to identify sets of neurons that implement the sketched morphology and connectivity pattern (**G1**).

**Spatial Motif Rendering (Sec. 7.2)**. After selecting particular motif instances, *MoMo* allows interactive 3D visualization of the respective neurons and their synapses and highlights the motif’s region of interest, such as highlighting the drawn segments in the visualization (**G3**).

## 6 Neuron Morphology-aware Motifs

Neuron morphology-aware motifs combine patterns related to neuron shape and their synaptic connectivity. In our proposed graph representation, a set of segments represents a single neuron (see Fig. 4b). Each segment is connected to neighboring segments or, through synaptic connections, to segments of another neuron. This contrasts previous connectome graphs where each neuron is represented as a single node connected by weighted edges indicating synapse count (see Fig. 4c).

**Fig. 4:**
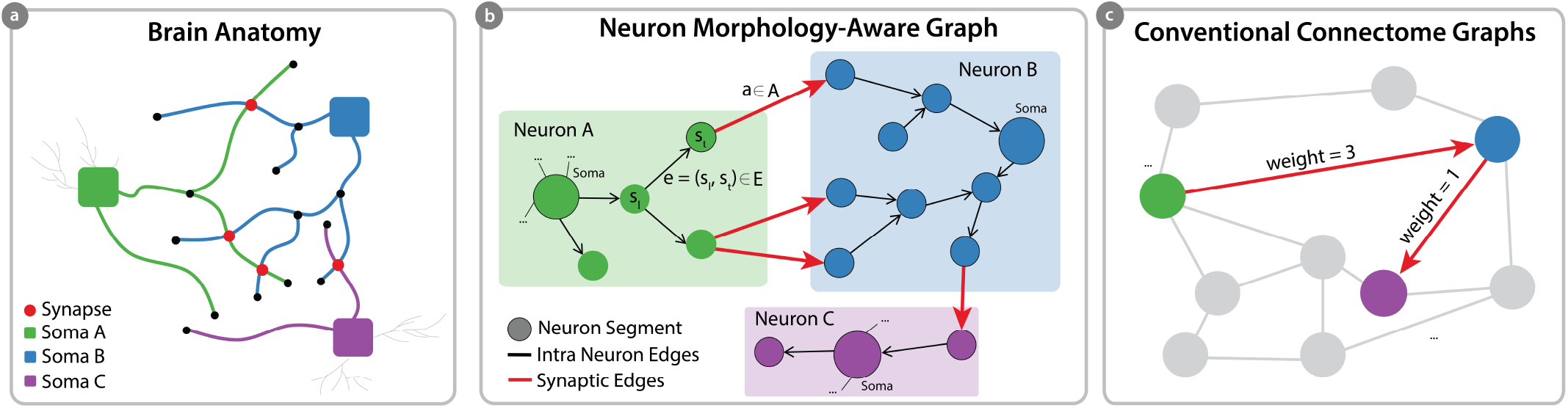
Neuron Morphology-Aware Graph. (a) Neurons build networks through complex tree-like arbors (dendrites/axons) that extend from each neuron’s soma. (b) Our graph representation captures the structure of wiring patterns by mapping each branch segment to a single graph node. Synaptic connections between segments are shown as red edges, while neighboring segments are connected through black edges. (c) Conventional connectome graph representations from previous work [39, 68] discard neuron shape information and fully abstract neurons into single graph nodes.

### 6.1 Formal Definition

Fundamentally, we construct a two-layer network, where one level describes *neuron connectivity* and the other layer *neuron morphology*. The connectivity layer abstracts the connectome as a directed graph *K* = (*N,C*), where a set of neurons *N* ={*n*_0_,…, *n*_*v*_} is connected through at set of synapses *C* = {*c*_0_,…, *c*_*u*_}. In the morphology layer, each tree-like neuron *n* ∈ *N* can be formalized as an acyclic skeleton graph *n* = (*V, E*). *V* is a set of 3D coordinates (vertices) connected through a set of edges *E* that describe the neuron’s centerline. Each synapse *c*_*ij*_ = {*p* ∈ ℝ^3^, (*i, j*)}∈*C* stores both a spatial position *p* and the indices (*i, j*) of the respective pre- and postsynaptic neurons. Here, we build a graph *G* that captures *both* neuron morphology *and* neuron connectivity. First, we partition each neuron skeleton *n* ∈ *N* into segments (Fig. 5). A segment *s* is a set of vertices *V*_*s*_ and edges *E*_*s*_ between two consecutive branching or terminal vertices. Formally, a segment *s* is defined as

**Fig. 5:**
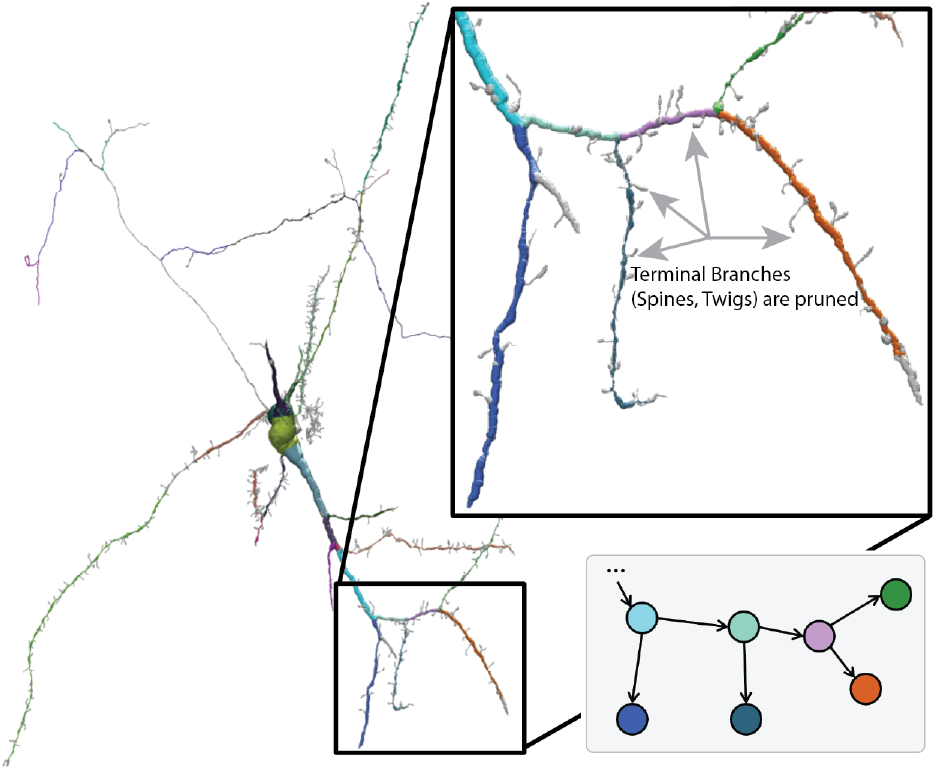
Neuron Segments. We partition each neuron into a set of segments. In our graph structure, each node represents a single segment. Neighboring segments are connected through edges. Terminal branches, such as spines or skeletonization artifacts/twigs, are being pruned during the data transformation. Data: Human temporal cortex / H01 [60].

**Fig. 6:**
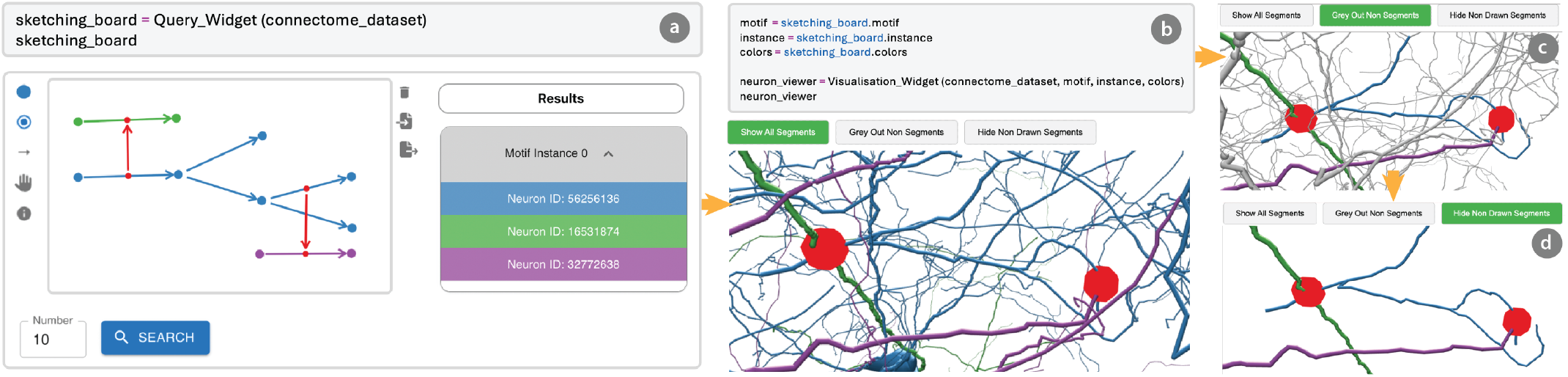
Interactive Motif Sketching & Interactive 3D Rendering. (a) Users start by sketching neuron segments and their synaptic connectivity using an interactive drawing board within a jupyter widget. Next, users can search for matching instances. A list of found motif instances is then displayed in the results panel. (b) Neurons of identified motif instances can be inspected in an interactive 3D rendering shown in a jupyter widget. Synapse locations are shown in red. (c) We allow users to grey out interactively or (d) hide neuron arbors not involved in the sketched motif to simplify creating correspondence with the motif sketch and the 3D spatial neuronal anatomy. Data: MICrONS [4].

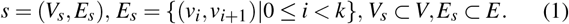

Additionally, all vertices but the start *v*_0_ and endpoint *v*_*k*_ must have degree 2. Each segment always consists of two or more vertices.

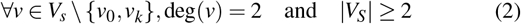

For each segment *s*, we also store a set of synapses *C*(*s*) that connect *s* to another neuron’s segment. Finally, we combine both layers in a neuron-morphology-aware graph *G* = (*S, E, A*). The vertices 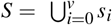 are the union of all previously computed segments. There are two types of edges. The first edge type *E* connects neighboring segments *s*_*l*_, *s*_*t*_ of the *same* neuron *n*_*m*_,

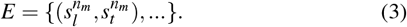

The second edge type *A* defines neuron connectivity and thus connects two segments of *different* neurons if they share synapses.

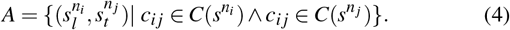

Next, given a motif *M*, we search for isomorphic subgraphs in *G*, such that

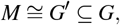

where *G*′ is an induced subgraph of *G. M* and *G*′ are isomorphic.

### 6.2 Data Transformation

Connectome data must be transformed into our graph representation. Skeletons are a standard and readily available format for representing 3D neuron data in connectomes. Skeletons store sample x,y,z coordinates and the radius along the centerline of a neuron. Transforming a set of skeletons and their respective synapses into our graph representation involves the following steps:

#### Pruning Terminal Branches

Neurons frequently contain numerous small terminal branches, such as spines or artifacts from the skeletonization procedure, that can obscure meaningful segment creation (see Fig.5). Thus, we iteratively prune terminal branches below a specified threshold *t*. Determining *t* depends on the properties of the datasets and is a hyperparameter of our graph representation. The goal is to suppress non-meaningful fine-scale morphology (e.g., spines or skeletonization noise) while preserving subcellular structures relevant for motif detection. As a rule of thumb, the pruning threshold shouldn’t exceed the average spine length to avoid losing meaningful structures. We evaluated thresholds ranging from 1–5µm on pilot datasets, assessing their impact on noise reduction and motif preservation. Based on empirical results and expert feedback, we selected dataset-specific pruning factors, with exact values reported in Section 9.

#### Synapse to Skeleton Mapping

Next, we establish a mapping between the location *p* of synapse and the closest skeleton vertex of its pre- and postsynaptic partner neuron. This mapping is necessary to determine which synapses correspond to which neuron segments.

#### Skeleton Downsampling

Next, we abstract the pruned neurons by focusing on more significant structural elements—the paths between branching points and endpoints. The exact curvature of these paths is disregarded to reduce computational complexity while maintaining essential connectivity. Thus, we downsample each neuron skeleton, so only branching points and endpoints remain. Branching- and endpoints are shown as black dots in Figure 4a. Those remaining points are a compact representation of the neuron’s arborization patterns.

#### Graph Assembly

The next step is restructuring the downsampled neurons into our neuron morphology-aware data representation. Neuron segments, defined as paths between branching points or endpoints, are represented as individual *nodes*. Neighboring segment nodes are connected through *intra neuron edges* (Fig 4b - black arrows). Synaptic connections between segments of different neurons become *inter neuron edges* types (Fig 4b - red arrows). Note that other properties such as neuronal compartmentalization (axon/dendrite) or synapse polarity (inhibitorty / excitatory) can be easily mapped to nodes and edges as well. The final graph can now be used to query morphological motifs. **Complexity**. The transformation from skeleton and synapse data to our graph representation scales linearly with dataset size. Let *N* be the number of neurons, *E* the average number of skeletal segments per neuron, and *S* the average number of synapses per neuron, giving a total time complexity of *𝒪*(*N* (*E* + *S*)). The pipeline includes loading skeletons, pruning small branches, downsampling, snapping synapses to skeleton nodes, and graph assembly. Neurons are processed independently, making the pipeline easily parallelizable. On standard hardware, we build a morphology-aware graph for 12,000 neurons and 500,000 synapses (FlyWire) in under 10 minutes on a 64-core, 256 GB RAM machine. For smaller datasets like MICrONS (1,700 neurons, 150,000 synapses), processing completes in under 90 seconds. Memory usage also scales linearly, driven mainly by connectivity and metadata storage.

## 7 Motif Visualization

In *MoMo*, visualizing neuron morphology-aware motifs is done through motif sketching and interactive 3D rendering of motif instances. We considered alternatives like visual subgraph selection, where users pick observed multisynaptic patterns directly from the visualization [3], and rule-based motif definitions using logical or structural constraints [39]. While visual selection is intuitive for highlighting known patterns, it relies on clean subgraph separation and does not scale well. Rule-based queries offer flexibility but are harder to express and computationally intensive. We chose sketch-based input for its speed, intuitiveness, and support for hypothesis-driven exploration.

### 7.1 Motif Sketching

The sketching board provides an interface for users to create motif sketches by defining neurons with distinct colors and specifying synaptic connections (Fig. 6a). Each node in the sketch represents either a branching point or an endpoint, while neuron segments are drawn in different colors selected from a menu, where each color corresponds to a distinct neuron. Users can add synaptic connections by selecting the appropriate tool and linking previously drawn segments (red edges). Both neuron segments and synaptic connections are directed, reflecting the natural flow of signals within and between neurons.

For example, Figure 6 illustrates a motif composed of three neurons (green, blue, and purple) and two synaptic connections. The *sketching board* accepts connectome datasets formatted in the neuron morphology-aware graph representation, allowing users to query the dataset based on the drawn motif. Matched motifs are dynamically displayed in a results list for exploration.

#### Querying for Morphological Motifs

Once the sketch is complete, users can initiate a query by pressing a designated button and specifying the desired number of results. The *MoMo* backend processes the request and returns a list of motifs matching the drawn example.

#### Reviewing Found Instances

Query results are displayed in a list view, where each detected motif instance is enumerated. Neuron IDs corresponding to the drawn segments are grouped and highlighted using the same colors as in the sketch for easy interpretation.

#### Sketch Reproduction and Sharing

Users can import and export motif sketches as JSON files, preserving segment information for future queries or sharing with collaborators, facilitating reproducibility and collaborative research.

### 7.2 Spatial Motif Rendering

*MoMo* provides an interactive 3D exploration of motif instances (see Fig. 6). After selecting a queried motif instance from the result list (Fig. 6a), users can visualize the corresponding neurons in 3D. The 3D viewer widget is launched via a Python function call (see Fig. 6b), enabling detailed inspection of the identified structures. *MoMo* provides three visualization modes, following a *focus & context* approach, to enable the user to correlate the sketched motif with the detailed 3D renderings of neurons from a specific motif instance (see Fig. 6b-d).

#### Full Instance Rendering

The first mode shows the full morphology of neurons involved in a motif instance (see Fig. 6b). This mode provides important spatial context. However, it can be hard to mentally map the sketched motif pattern to the 3D neuron rendering due to the cluttered neuron morphologies.

#### Focused Instance Rendering (Color)

In the second mode, we defocus neuronal branches that are unrelated to the sketched motif by rendering them in gray (see Fig. 6c). This helps to identify the neural processes involved in the motif easily visually but can still be perceived as cluttered due to occlusion effects.

#### Focused Instance Rendering (Pruning)

The third mode prunes unrelated neuron branches that are unrelated to the motif. This enables clear inspection of the neuronal segments that implement a motif pattern (see Fig. 6d). However, this view lacks important spatial context and is thus complemented by the previous two visualization modes.

In all modes, synapses are represented as red spheres within the visualization. The user can switch between those three modes interactively in the user interface (see Fig. 6b-d).

## 8 Interactive Motif Queries

Our neuron morphology-aware graph representation extends conventional connectome graphs by introducing additional nodes and edges, increasing complexity significantly. In the MICrONS dataset, 1,712 proofread neurons connect via roughly 150,945 synaptic edges. Transformed into our morphology-aware graph, this yields over 390,000 nodes and 400,000 edges.

### Computational Challenges

Isomorphic subgraph search is an *NP-complete* problem [13], with worst-case complexity growing exponentially with graph size. While manageable for small graphs, our large-scale representation leads to prohibitively long runtimes even for small motifs. Such delays hinder interactive use cases like *MoMo*, where users expect near-instantaneous motif sketching and instance inspection.

### Parallel Motif Discovery

To enable rapid exploration, we integrated the parallel VF2-PS [15] algorithm into *MoMo*. VF2-PS reformulates subgraph isomorphism as a state-space search [48], where each state represents a partial or full mapping between motif graph *M* and host graph *G* vertices. This approach supports efficient parallelization across many cores. For instance, VF2-PS can reduce query times for simple motifs by up to 48 × compared to conventional NetworkX-based queries (see Supplementary Table 1).

### Custom Motif Discovery Features

We extended the VF2-PS algorithm with three custom features to enhance interactive motif analysis.

First, *wildcards* (Fig. 7) allow users to leave certain neuronal morphology attributes undefined, enabling exploration of structural hypotheses without requiring complete morphology knowledge. For example, to examine whether multiple synaptic connections between two neurons occur in distinct spatial regions, users can sketch disconnected neuron segments in different colors and specify synapses accordingly. Second, *color matching* ensures nodes correspond to distinct neurons and their connections. We introduced a post-processing step that tracks colors assigned to nodes and edges; each motif instance is validated to confirm its color pattern matches the input subgraph, meaning colors map to unique neuron IDs. Third, users can set *query limits* to bound the number of motif instances returned (Fig. 6a), allowing interactive exploration without enumerating all instances in the full dataset.

**Fig. 7:**
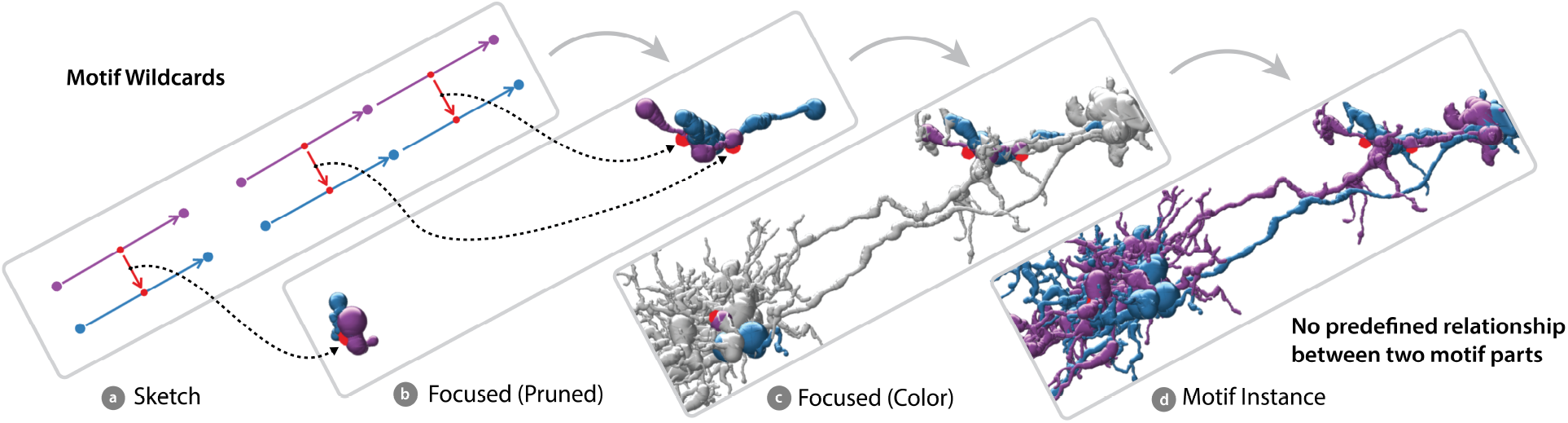
Wildcard Feature. (a) *MoMo* enables users to leave specific parts of the motif query undefined. (b) Pruned and focused view of the retrieved instance, placed adjacent to the sketch to enable direct visual comparison. Black dashed arrows link sketched synapses (a) to their corresponding synapses in the instance (b). (c) Focus-and-context rendering: unrelated neuronal branches are grayed out to highlight relevant morphology. (d) Full, unpruned view of the retrieved instance for reference. Data: FlyWire [16].

## 9 Data and Implementation

*MoMo* requires a connectivity network and proofread reconstructions of neurons represented as 3D skeletons and synapses, along with their spatial locations. *MoMo* then processes the input data and transforms it into the neuron morphology-aware graph representation. Once the data is transformed, all subsequent computations can be executed during runtime. We tested *MoMo* on two different connectome datasets.

### FlyWire Data

We tested *MoMo* on a subset of the FlyWire connectome [16] that includes neurons from the *medulla intrinsic* (Mi) and T4 families, *Lamina Intrinsic, Lobula Intrinsic, Centrifugal*, and *Distal Medulla* neurons. All these neurons are located in the right optical lobe, bringing the total to 12,803 neurons with 532,530 synaptic connections. The final neuron morphology-aware graph has 956,729 nodes and 654,778 intra-neuron edges. Based on experimental evaluations and expert feedback, we set the pruning factor to 3 µm for this dataset.

### MICrONS Data

As a second dataset, we tested *MoMo* on a set of 1,712 proofread neurons from the MICrONS dataset [4]. This data describes the detailed synaptic wiring of a mouse’s primary visual cortex. The respective neuron morphology-aware graph contains 392,438 nodes, 250,923 intra-neuron edges, and 156,945 synaptic edges. For this dataset, we applied a pruning factor of 1 µm, following the same evaluation process as for the Flywire dataset.

### Implementation

*MoMo* is a modular Python library designed for Jupyter environments. Its motif sketching interface (Fig.6a) and 3D visualization widget (Fig.6b–d) are separate components, each embedded in a Jupyter cell. The system leverages the *AnyWidget* framework [37] to integrate custom JavaScript components. Both widgets are built with JavaScript/React, while data processing runs in Python. *MoMo* builds on libraries including *Navis* [58] for neuron data handling and *Paper*.*js* for vector graphics in motif drawing. The 3D skeleton rendering uses *Three*.*js* with SharkViewer [71]. The *VF2-PS* algorithm is implemented in Arachne [2, 18], providing access to parallel property graph structures [54]. Our code and example scripts for transforming skeletons into our graph representation are publicly available on GitHub (see abstract). Datasets will be released upon paper acceptance.

## 10 Evaluation

We evaluate *MoMo* through a series of case studies, assessing the usefulness for real-world neuroscience research questions.

### Participants

We evaluated *MoMo* with four experts (**P1–P4**): three male and one female, from Harvard University, Janelia Research Campus, the University of Würzburg, and Freie Universität Berlin. The group comprised one professor, one research scientist, and two post-doctoral researchers, all specializing in the analysis of neuronal circuits reconstructed from EM image data. Additionally, two participants had prior experience using graph abstraction tools for connectome analysis. Their diverse expertise provided valuable neuroscientific perspectives, reinforcing the tool’s broad applicability.

### Session Structure

Each session lasted about an hour, conducted either in person or via interactive Zoom with remote screen control. Except for the first participant (a pilot study during tool development), all experts engaged hands-on with *MoMo*. The pilot participant provided early feedback and use cases that shaped the tool’s design. Sessions began with an introduction to *MoMo*, highlighting its capabilities and novel integration of neuronal morphology in motif analysis. Participants then sketched example motifs and analyzed retrieved instances, followed by discussions on biological relevance, applications, and areas for improvement. Each case study describes the expert’s analysis goal, biological context, and how *MoMo* supported their investigation. Three of four focused on the relationship between neuron morphology and synapse polarity (inhibition/excitation). While *MoMo* does not explicitly store polarity, experts agreed that understanding the underlying connectivity is a critical first step toward analyzing these phenomena. The tool enables exploratory analysis of motif instances that *could* support relevant neural computations. Finally, we synthesize insights across all case studies, highlighting strengths, challenges, and future improvement opportunities.

### 10.1 Pilot Study: Investigating Shunting Inhibition

As an initial pilot study, we collaborated with **P1** to explore potential applications of *MoMo* during its development. They suggested investigating the biologically relevant use case of *shunting inhibition*. This phenomenon exemplifies a case where both neuronal morphology and connectivity must be considered, making it challenging to study with existing graph-based tools.

#### Understanding Shunting Inhibition

*Shunting inhibition* is a neural mechanism in which inhibitory inputs suppress excitatory signals, preventing their propagation. This occurs when an inhibitory neuron synapses onto a neuron receiving an excitatory input, but the inhibitory input is positioned upstream relative to the excitatory one. As a result, the inhibitory signal can cancel or *shunt* the excitatory effect before it travels further through the neuron [32, 44].

#### Applying *MoMo* to Shunting Inhibition

During the remote session conducted via Zoom, the expert instructed us to sketch multiple motifs representing possible instances of shunting inhibition. One of the sketched motifs can be seen in Fig. 8a. The sketched motif comprises a network of three interconnected neurons: a blue neuron with four horizontally connected segments, a green neuron segment linked to its leftmost part, and a purple neuron segment synapsing onto its rightmost part. Given the left-to-right signal flow, the expert hypothesized that an inhibitory signal from the purple neuron could shunt an excitatory input from the green neuron, preventing its transmission along the blue neuron. The expert also instructed us to iteratively create additional motifs with varying numbers of blue segments to explore how the tool would handle different configurations. Following the motif query, they explored multiple matching instances within the Flywire dataset [16]. The expert examined the visualized results, assessing whether the retrieved structures aligned with the desired morphological and connectivity patterns. Figure 8 illustrates one of the identified instances across different view modes in *MoMo*. In Figure 8c, only the sketched segments are highlighted in their respective colors, with the rest of the neuron structures grayed out for clarity. Figure 8e shows the four segments of the blue neuron are colored differently to emphasize different segments of the blue neuron in the motif sketch.

**Fig. 8:**
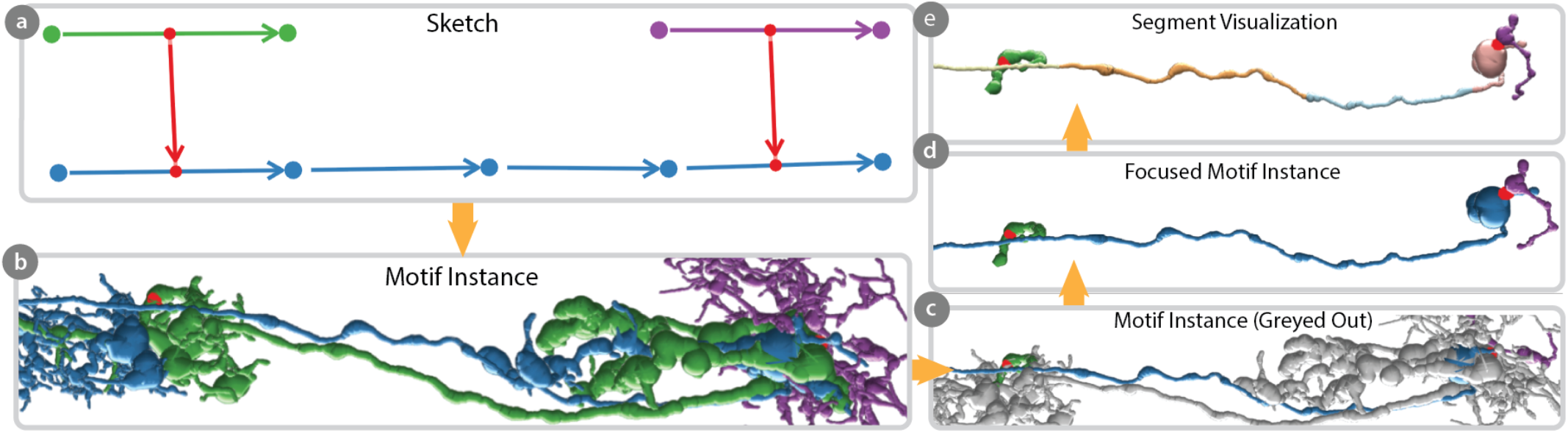
Pilot Study: Query for Potential Shunting Inhibition. (a) The sketch illustrates a three-neuron motif (green, blue, and purple). The green neuron connects to the leftmost segment of the blue neuron, while the purple neuron connects to the rightmost segment. Thus, assuming the purple neuron provides inhibitory input, it could suppress the excitatory input from the green neuron due to the left-to-right signal propagation within the blue neuron. (b-d) The renderings show an exemplary motif instance queried with *MoMo* at different levels of focus. (e) We show the four corresponding segments of the blue neuron by assigning each segment a distinct color. Data: FlyWire [16].

### 10.2 Case Study 1: Lateral Inhibition

To further assess *MoMo*’s applicability, we conducted a case study with **P2**. They highlighted *MoMo* ‘s potential as a complementary approach to tools like Vimo that abstract entire neurons as nodes. In their view, Vimo is well-suited for initially defining broader connectivity constraints, while *MoMo* becomes particularly valuable when morphology needs to be considered alongside connectivity for more detailed investigations. They suggested using the tool to investigate *lateral inhibition*. Unlike the pilot study, this expert directly interacted with *MoMo* in person, allowing for more hands-on evaluation of the tool’s query and visualization capabilities.

#### Understanding Lateral Inhibition

*Lateral inhibition* is a fundamental neural process where inhibitory neurons suppress the activity of neighboring excitatory neurons, enhancing contrast in sensory processing and aiding pattern recognition. This typically involves inhibitory synapses onto adjacent excitatory neurons, refining signal transmission by suppressing their responses. A classic example (Fig. 9a) shows Neuron A sending excitatory signals to Neuron B, which then inhibits neighboring neurons C and D. This prevents lateral spread of action potentials and sharpens the signal contrast. Such mechanisms enable the distinction of stimuli by enhancing the excited neuron’s response while dampening its neighbors. Identifying these motifs in large-scale connectomic data requires tools that analyze both synaptic connectivity and spatial relationships between neurons.

**Fig. 9:**
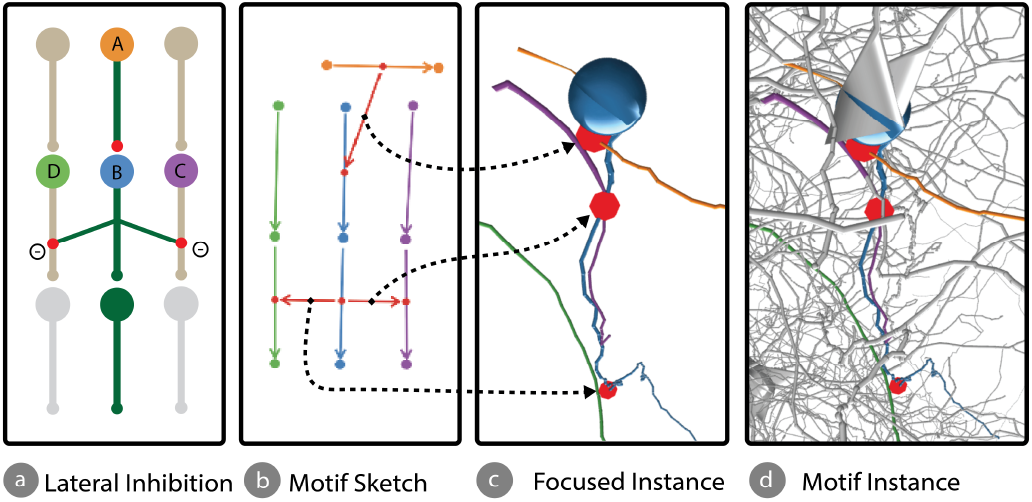
Case Study 1. Lateral Inhibition. (a) Schematic representation of *lateral inhibition*. (b) User-drawn sketch illustrating a four-neuron example with connectivity and morphology matching the schematic. (c, d) Visualizations of a retrieved motif instance at different levels of focus. Black dashed arrows link synapses in the sketch (b) to their corresponding synapses in the retrieved instance (c). Data: MICrONS [4].

#### Applying *MoMo* to Lateral Inhibition

In this case study, the expert used *MoMo*’s sketching interface to draw multiple motifs corresponding to potential instances of *lateral inhibition*. These motifs typically involved central neurons receiving excitatory inputs from neighboring neurons and subsequently potentially inhibiting the activity of these neighboring neurons. Some drawn motifs matched existing structures in the dataset, while others did not. After each motif sketch, the expert utilized *MoMo*’s visualization interface to inspect the identified instances and assess whether they corresponded to the expected anatomical and connectivity patterns of *lateral inhibition*. This process was iterative—after reviewing the results, the expert modified the sketch and conducted another motif search to refine the analysis. The expert highlighted that *MoMo*’s ability to incorporate detailed morphological structure greatly facilitated the investigation of complex phenomena like *lateral inhibition*, enabling a more nuanced exploration than possible with purely connectivity-based tools. Figure 9b presents an example of a potential *lateral inhibition* motif, derived from the expert’s sketched motifs and follow-up discussions. In this case, the orange neuron is depicted as potentially making an excitatory synaptic connection onto the blue neuron, which then forms potential inhibitory synapses onto the neighboring purple and green neurons, dampening their responses and creating a feedback inhibition loop. Figure 9c provides a simplified view of an identified instance from the MICrONS dataset [4], emphasizing only the relevant neuron segments and synaptic connections from the original sketch. In contrast, Figure 9d presents the same instance with the full neurons in grayscale, preserving the sketch’s color to provide broader structural context.

### 10.3 Case Study 2: Feed Forward Excitation

To assess *MoMo* ‘s usability for researchers who are not familiar with graph-based representations of neuronal data, we conducted a case study with **P3**, an expert specializing in the detailed analysis of physical neuronal structures, such as images and 3D reconstructions, rather than relying on graph abstractions. This expert’s perspective was invaluable in understanding how *MoMo* could enhance the study of neuronal morphology using graph-based tools. For this case study, **P3** chose to investigate *feed-forward excitation* and the presence of dense synaptic connections within neurons.

#### Biological Significance of Feed-Forward Excitation

*Feed-forward excitation* allows a neuron to propagate excitatory signals downstream, facilitating rapid signal transmission and integration across brain regions. It often involves both convergence—where multiple presynaptic neurons excite a single postsynaptic target—and divergence—where one presynaptic neuron excites multiple targets (see Supplementary Fig. 12). This organization enhances feature selectivity and improves the signal-to-noise ratio in sensory and motor pathways [73]. Dense synaptic connections occur in regions with a high concentration of synapses, facilitating complex neural processing. These connections are crucial for brain areas involved in learning, memory, and sensory processing, supporting the integration of large amounts of information for tasks like pattern recognition and cognition.

#### Applying *MoMo* to Feed-Forward Excitation

During the Zoom session, the expert engaged with *MoMo*’s sketching interface, using remote control to explore motifs related to *feed-forward excitation* and dense synaptic connections. This was the expert’s first time considering a graph-based abstraction of the connectome data, which required a shift in perspective from their typical approach of analyzing physical neuronal structures. They actively explored the found instances of the drawn motifs by leveraging *MoMo* ‘s various view modes, rotating and zooming in and out of the visualized motifs. The real-time querying feature was particularly advantageous, allowing the expert to refine and visualize their hypotheses iteratively. This dynamic exploration facilitated a deeper understanding of how complex neural circuits are connected and how spatial arrangements of neuronal structures influence connectivity patterns. The ability to interactively adjust the visual representation helped bridge the gap between abstract graph-based data and the expert’s more familiar visualization of neuronal morphology.

### 10.4 Case Study 3: Center-Surround Receptive Fields

This case study was conducted in collaboration with **P4**, who was already familiar with previous tools that utilized a sketch-based graph abstraction for connectomics data. After discussing several initial ideas, the expert chose to investigate the phenomenon of *center-surround receptive fields* for this session.

#### Understanding Center-Surround Receptive Fields

In sensory neuroscience, neurons process information from their surroundings in structured patterns. A common motif in visual processing is the *center-surround receptive field*, where a neuron’s response depends on the contrast between the center and the surrounding region of its input. This structure enhances edge detection and contrast sensitivity, which are critical for vision. Neurons tuned to this pattern fire most strongly when a bright stimulus is in the center and a darker region surrounds it (or vice versa). This mechanism is fundamental in biological vision systems, including those in flies and mammals, and is analogous to computational filters like *Gabor filters* used in artificial vision models.

#### Applying *MoMo* to Center-Surround Receptive Fields

During the Zoom session, the expert used *MoMo* ‘s sketching and visualization interface via remote control to analyze and refine neuron motifs relevant to *center-surround receptive fields*. After several iterations, they finalized the sketch shown in Figure 10a. The green neuron represents the hypothesized central excitatory unit, responsive to bright stimuli at the receptive field’s center. The blue neuron acts as a localized feature detector, selectively responding to smaller bright regions to enhance fine detail detection. Surrounding these, the purple and orange neurons represent potential surround inhibitory units, activated by darker areas around the center to support contrast detection. Using *MoMo* ‘s real-time query feature, the expert identified candidate instances of the *center-surround receptive field* phenomenon within the MICrONS dataset [4] (see various views of one instance in Figures 10b and c). They explored these motifs through *MoMo* ‘s visualization interface, leveraging its multiple views to examine morphological and connectivity patterns consistent with the expected characteristics.

**Fig. 10:**
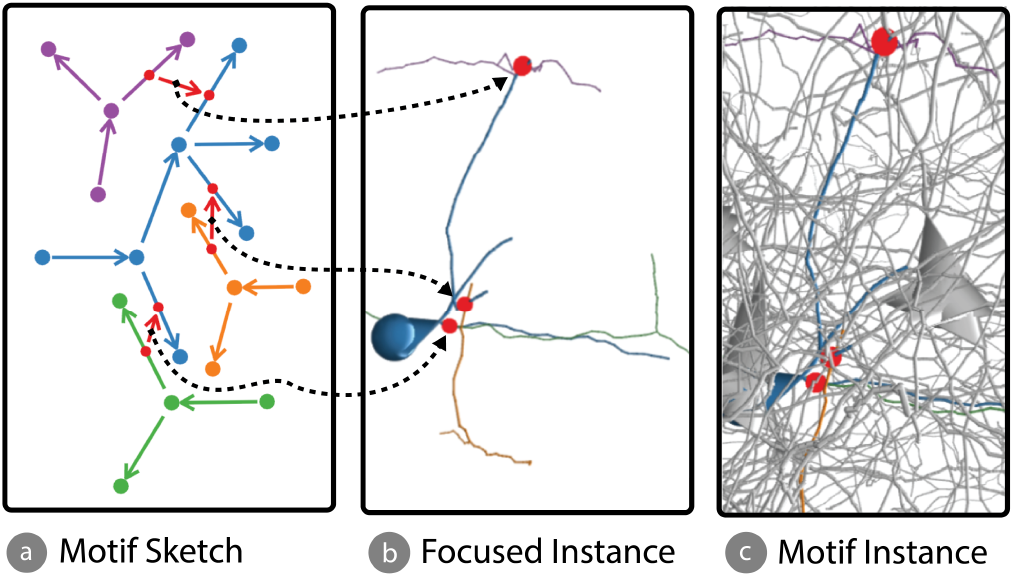
Case Study 3: Center-Surround Receptive Fields. (a) Motif sketch, which could correspond to a potential *center-surround receptive field*. Key components include a green neuron (potentially the central excitatory unit), a blue neuron (potentially the localized feature detector), and purple and orange neurons (potentially the surround inhibitory neurons). (b, c) Renderings of identified instances of the motif, queried using *MoMo*, shown at different levels of focus. Black dashed arrows link synapses in the sketch (a) to their corresponding synapses in the retrieved instance (b). Data: MICrONS [4].

### 10.5 Key Evaluation Findings

*MoMo* effectively fulfills its core goals and tasks outlined in Section 4 by offering an interactive, morphology-aware motif analysis tool for connectomics research. Across all case studies, experts highlighted its intuitive interface, real-time querying, and seamless integration of sketching and visualization. However, while the tool offers several advantages, certain limitations and areas for improvement were identified, which are discussed below.

#### Efficient and Intuitive Motif Identification

One of *MoMo* ‘s most valued strengths is its ability to facilitate rapid motif identification (**G1, T1**) through an interactive sketching interface (**G2, T2**). Experts particularly appreciated the tool’s capacity to refine and adjust queries in real time, especially when no exact matches were initially found, like in the first case study. This flexibility lowers the barrier to entry for researchers unfamiliar with complex query languages (**G4, T4**). However, relying on user-generated sketches introduces subjectivity, as motif definitions may vary across users. Future enhancements could include predefined motif templates or automated suggestions.

#### Enhancing Connectivity and Morphology Analysis

*MoMo* effectively supports combined connectivity and morphology queries, enabling detailed circuit-level analyses (**G2**). This was demonstrated in studies on *shunting* and *lateral inhibition*, where experts specified spatial arrangements of potential excitatory and inhibitory synapses to retrieve biologically relevant instances. However, search accuracy depends on preprocessing: pruning fine structures improves efficiency but risks losing critical neuronal details. Selecting the right pruning factor is essential to balance speed and fidelity. Experts also recommended adding semantic information—such as branch lengths or region-specific queries—to further enhance *MoMo* ‘s utility.

#### Insect vs. Mammalian Brains

We tested *MoMo* on both the *FlyWire* and *MICrONS* datasets, allowing participants to switch between them. This raised the question of whether our abstraction approach suits certain connectomes better. Our segment-level representation aligns more naturally with mammalian neurons, where terminal branches (twigs) are relatively uniform and consistently pruned. In contrast, insect neurons (*FlyWire*) show greater morphological variability, requiring dataset-specific pruning adjustments. Fig.11a shows a mammalian neuron with small, uniform twigs, while Fig.11b highlights the complexity of insect neurons, especially in the yellow-circled region. While preprocessing is simpler for mammalian neurons, *MoMo* remains adaptable to insect datasets through parameter tuning.

**Fig. 11:**
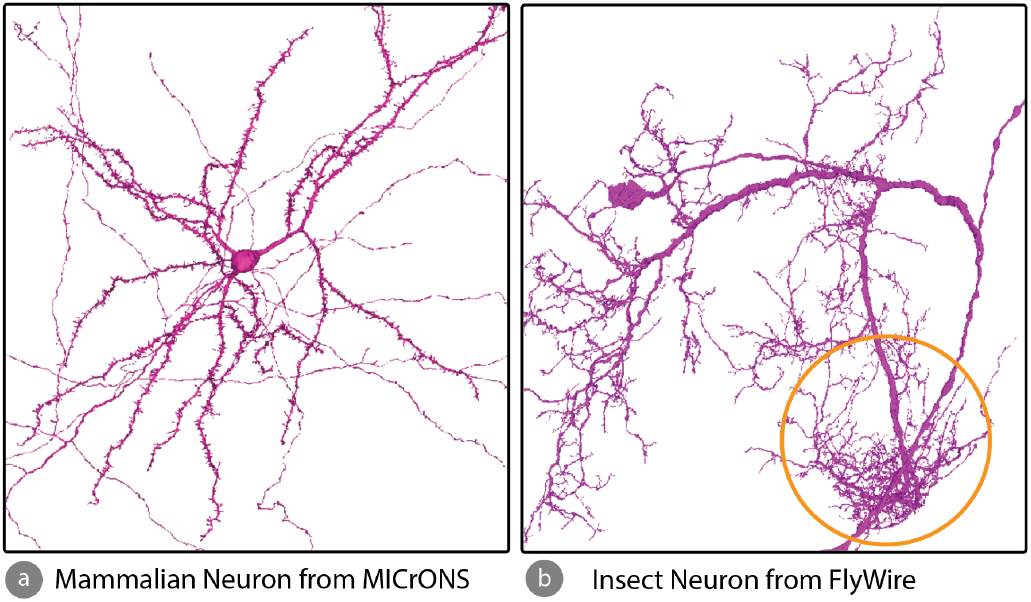
Mammalian Neurons vs. Insect Neurons. Comparison of neuronal structures between the *MICrONS* and *Flywire* datasets. (a) A mammalian neuron from *MICrONS*, characterized by small, uniform terminal branches. (b) An insect neuron from *Flywire*, displaying more intricate and variable terminal branches, particularly in the yellow-circled region. Thus, mammalian neurons are overall better suited for *MoMo*.s

#### Improving Spatial Analysis and Visualization

*MoMo* ‘s ability to correlate motif queries with 3D neuron reconstructions (**G3, T3**) is a significant advantage. Experts appreciated the tool’s multiple visualization modes, which aid spatial reasoning about synaptic interactions. However, some participants found interpreting dense 3D structures challenging. Future improvements could include occlusion-aware highlighting or interactive slicing to enhance clarity in highly interconnected regions. Another key challenge lies in the wildcard feature, which allows flexible matching of neuronal segments. While this feature provides adaptability, it currently lacks the ability to constrain wildcard segments to specific neuronal regions. For example, in the *feed-forward excitation* case study, a constraint ensuring that synaptic connections occurred on a particular neuron arbor would have improved specificity. Introducing constraint options for wildcards could enhance motif refinement while preserving flexibility.

#### Limitations of Motif Querying

While *MoMo* enables segment-level motif sketching and structural queries, some limitations remain. Users cannot specify exact segment lengths or precise synapse positions along a segment, as sub-segment resolution is not supported. Quantitative constraints (e.g., “at least three inputs”) are also unavailable, limiting expressiveness for some patterns. While effective for small to moderately sized motifs (typically 3–4 neurons with 3–6 segments and synapses), querying larger or more complex motifs is challenging due to combinatorial search space and sketching usability. As noted by **P2**, *MoMo* works best alongside higher-level tools better suited for motifs with 5+ neurons.

### 11. Conclusion and Future Work

In this paper, we introduced a novel method for interactively searching and analyzing neuron morphology-aware motifs in large connectome graphs. Our approach enables exploration of complex neural structures and their functional relationships across datasets like *FlyWire* and *MICrONS*. By focusing on segment-level connectivity patterns, it facilitates identifying motifs—such as lateral inhibition—that were previously difficult to query. Through case studies, we demonstrated the tool’s effectiveness in supporting exploratory analysis and providing insights into neuronal function. We also presented a fully functional prototype enabling efficient, interactive analyses. Looking ahead, several exciting directions remain. First, comparative visualization methods are needed to correlate morphology-aware connectivity across specimens of different sexes or developmental stages, which poses challenges such as establishing visual correspondences. Second, *MoMo* currently supports morphological but not spatial motif queries—patterns involving spatial coordinates alongside morphology, like helix motifs. Integrating spatial queries would require additional data and increase computational demands. Third, alternative motif definition interactions, such as direct selection in 3D renderings or scribbling-based inputs, should be explored. Finally, beyond hypothesis-driven motif searches, integrating graph-based machine learning with interactive visualization could reveal unknown motifs; a key challenge will be effectively visualizing these automatically identified morphological patterns.

## Acknowledgements

We thank the Chapel and Arkouda communities for their guidance and support. This research was supported in part by the NSF grants NCS-FO-2124179, CCF-2109988, OAC-2402560, and CCF-2453324.

