## Supplement Table 1 and Supplement Figure 12 for "MoMo - Combining Neuron Morphology and Connectivity for Interactive Motif Analysis in Connectomes"

Table 1: **Algorithmic Evaluation.** Execution time comparison for motif querying between Arachne and NetworkX. Arachne is utilized in *MoMo*. Performance measured on a system with two AMD EPYC 7713 CPUs (64 cores each) and 1TB RAM. Data: FlyWire.

| Subgraph | VF2-PS (sec) | NetworkX (sec) | # Instances |
|---|---|---|---|
| 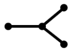 | **2.48** | 336.45 | 696,460 |
| 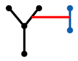 | **3.62** | 173.75 | 191,690 |
| 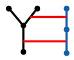 | **2.88** | 5,980.54 | 5,048 |
| 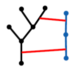 | **339.46** | 16,436.85 | 2,308 |
| 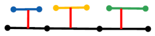 | **1.56** | 435.07 | 44,657 |
| 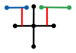 | **78.77** | 810.23 | 161,842 |
| 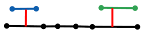 | **4.10** | 1,018.23 | 179,255 |
| 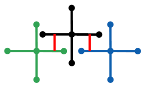 | **38.06** | >12,000 | 4,992 |

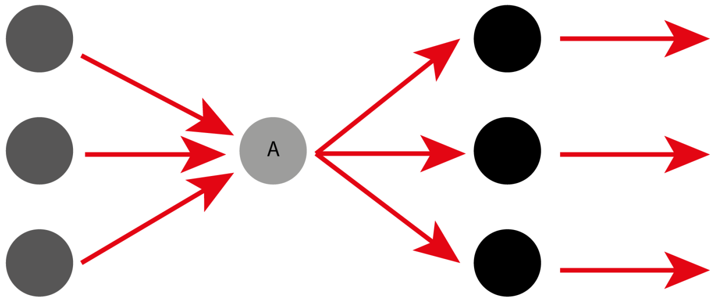


Fig. 12: **Feedforward Excitation** Illustration of convergent and divergent feed-forward excitation: multiple upstream neurons converge onto a single target neuron (A), which then diverges to excite multiple downstream neurons. The figure was adapted from Zheng, S. et al [73]
